# Neanderthal Genomics Suggests a Pleistocene Time Frame for the First Epidemiologic Transition

**DOI:** 10.1101/017343

**Authors:** Charlotte J. Houldcroft, Simon J. Underdown

## Abstract

High quality Altai Neanderthal and Denisovan genomes are revealing which regions of archaic hominin DNA have persisted in the modern human genome. A number of these regions are associated with response to infection and immunity, with a suggestion that derived Neanderthal alleles found in modern Europeans and East Asians may be associated with autoimmunity. Independent sources of DNA-based evidence allow a re-evaluation of the nature and timing of the first epidemiologic transition. By combining skeletal, archaeological and genetic evidence we question whether the first epidemiologic transition in Eurasia was as tightly tied to the onset of the Holocene as has previously been assumed. There clear evidence to suggest that this transition began before the appearance of agriculture and occurred over a timescale of tens of thousands of years. The transfer of pathogens between human species may also have played a role in the extinction of the Neanderthals.

## Introduction

Current models of infectious disease in the Pleistocene tell us little about the pathogens that would have infected Neanderthals (*Homo neanderthalensis*). If we consider the work of Cockburn^1,2^, Omran^3^, and Barrett^4^, who argue that infectious disease only started to seriously impact human groups after the development of agriculture during the Holocene, we must assume that Neanderthals lived in a time and place largely free of acute infectious disease. The current epidemiologic transition model associates most infectious diseases that cause significant mortality with changing living conditions connected with the rise of agriculture, increased sedentism and higher population densities. As Pleistocene hunter-gatherers, Neanderthals should not be at risk from the “pestilences” of Omran’s thesis. However, new genetic evidence has the potential to change our view of Neanderthal pathology, and perhaps even the model of the first epidemiologic transition.

Firstly, we must consider the current tools for studying infectious disease in the Pleistocene period. Before the advent of ancient DNA sequencing methods, researchers were limited to studying the skeletal pathologies of humans and Neanderthals from this period (fossilised evidence of bones responding to infection and inflammation). However, only a limited subset of infectious diseases leaves behind these lesions. The publication of high-quality Neanderthal and Denisovan genomes gives us a new opportunity to study Pleistocene infectious disease. Researchers from a range of disciplines interested in the evolution of the modern suite of infectious disease can also draw inferences from this new source of data. As a result of making comparisons between modern humans genomes, seeking genetic polymorphisms which vary in function or frequency between populations, and by also comparing human genomes with high-quality Denisovan and Neanderthal genomes, we are beginning to find evidence of introgressed Neanderthal and Denisovan alleles and haplotypes which have functions in immunity and the response to infection^5–7^. The persistence of these regions of DNA in some modern human genomes suggests they may have conveyed a selective advantage, increasing the fitness of anatomically modern humans (AMH) when dispersing into new environments. Through comparisons of genetic data with skeletal evidence of infection, it is increasingly to analyse which pathogens shaped the evolution of modern humans and their closest relatives. Furthermore, ancient DNA technology now encompasses pathogen DNA, and in the future it may be possible to sequence pathogen DNA directly from Neanderthal remains – including pathogens that do not cause skeletal lesions. The genomes of extant human pathogens are shedding light on the antiquity of these infections in hominins.

We will discuss the evidence for infectious disease in Neanderthals, beginning with that of infection-related skeletal pathologies in the archaeological record, and then consider the role of infection in hominin evolution. We have a synthesised current thinking on the chronology of emergence of notable European disease packages (Table 1). Finally, we will consider what implications this evidence has for the classical model of the first epidemiologic transition.

## The Neanderthal Fossil Record

*Homo neanderthalensis* was a large bodied hominin that inhabited Eurasia widely from approximately 250,000 to 28,000 years ago^8^. Neanderthals occupied a hunter-gatherer subsistence niche, forming small bands of approximately 15-30 individuals^8^. Archaeological analysis suggests that while Neanderthal groups were relatively self-sufficient there was some level of exchange and transfer of materials^9^ The Neanderthal fossil record of some 400 individuals represents one of the largest collections of extinct hominin remains and is larger than that of contemporary Pleistocene *Homo sapiens* fossils. Genetic estimates for Neanderthal population size vary but agree that total numbers were small. Most studies suggest that the effective population size was between 3000-25000 (peeking around 50KYA before gradual decline)^10–14^. Bocquet-Appel & Degioanni^15^ have suggested that the actual population could have theoretically reached 70,000 but agree that the effective population size would have been much smaller due to the impact of environmental and ecological pressures. Thus, the fossil record presents researchers with a broad sample of the whole population. Despite the breadth of skeletal material and the large number of pathological lesions described, the Neanderthals are still viewed as existing in a hunter-gatherer epidemiologic paradigm, an effect of the traditional approach of describing each fossil in relative isolation. Conversely, while systematic population level studies have shown that the Neanderthals sustained high levels of traumatic injury^16,17^, the same methods have largely not been applied to infectious disease. When reviewed as a species there is evidence that along with traumatic injury the Neanderthals displayed a broad range of dental pathology and degenerative diseases as well as a large amount of non-specific infection^18–21^.

From the perspective of the current epidemiologic transition model, the Neanderthals’ small group size and limited exchange networks suggests that they could not act as reservoirs for infectious diseases. Also, much has been made of the Neanderthals’ apparent lack of cognitive and technological sophistication^22^. There is no logical reason to suppose infectious diseases were unknown to Neanderthal groups based on the fossil evidence. Indeed, the structure of Neanderthal groups would have made disease a potent factor in any demographic collapse related to extinction events^23^. As our understanding of Neanderthal biology and behaviour becomes more sophisticated we are presented with a hominin which was arguably every bit as intelligent and adaptive as *Homo sapiens*^24–26^. Their extinction and our survival potentially questions any innate superiority in *Homo sapiens*. Recent genetic analysis that suggests interbreeding further calls into question the real nature of the divisions traditionally drawn between the two alpha human hominins.

## Innate, adaptive and archaic immunity in hominin genomes

2010 saw the publication of the draft Neanderthal genome sequence^27^, which revealed that humans living outside Africa have a small proportion of Neanderthal ancestry – ̃2% of their genome^28^. Three Neanderthal genome sequences are available: a draft sequence from Vindija in Croatia, the composite sequence of DNA from bones from three different layers (inferred to be from different individuals), dating from between 38-45kya^27^; a low-coverage sequence of a Neanderthal found in Mezmaiskaya in the Caucasus, from a layer dated as 60-70kya; and a high-quality Neanderthal genome from the Altai region^5^, dated to 29-45kya. The data set is growing constantly, recently bolstered by a 49kya Neanderthal exome sequence (the ̃1% of the genome which codes for proteins) from El Sidron in Spain, and a further 44kya exome from Neanderthal remains recovered from Vindija^29^. Comparisons of these genome and exome sequences (although taken from only a handful of individuals) to those of modern humans have identified several regions of genetic similarity between humans and Neanderthals that are thought to have arisen from admixture between these two hominins. Approaches to identifying introgressed Neanderthal regions in the human genome which may be adaptive have looked for a range of different kinds of variation, from haplotype blocks hundreds of kilobases long, to single nucleotide polymorphisms (SNPs).

One such putatively introgressed region plays a role in innate immunity to viral infections. A haplotype containing *OAS1*, *OAS2*, *OAS3* of Neanderthal origin has been found in some modern human genomes^30^. These genes activate RNase L to degrade viral RNA. One Neanderthal derived SNP in *OAS2*, rs15895, is associated with response to tick-borne encephalitis virus disease in Europeans^31^. This is a disease found in forested areas of northern, central and eastern Europe, which would have formed a major part the Neanderthals’ typical ecosystem^8,32^. Did this pathogen represent a particular selection pressure for AMH colonising Europe, unlike genetically adapted Neanderthals?

There is also evidence for Neanderthals contributing to the innate immune system in modern Papua New Guinea. A study by Mendez^30^ found a haplotype carrying three genes (*STAT2*, *ERBB3*, *ESYT1*) to be absent in Africans, but present at variable frequencies outside Africa, peaking at 54% in Melanesians. The divergence time for the putatively introgressed Melanesian *STAT2* haplotype and the Neanderthal *STAT2* haplotype is 78kya, compared to a divergence time between the Human Reference Sequence haplotype and Neanderthal haplotype of 609kya. *STAT2* is involved in the interferon-alpha response to viral infections, including dengue^33^, influenza and measles ^34^. It is of note that *STAT2* interacts with other putatively introgressed Neanderthal genes, discussed above: *OAS1*-*3*.

Sankararaman and colleagues^7^ scanned the genomes of modern Europeans and Asians for evidence of individual SNPs that have introgressed from Neanderthals, a number of which have been associated with immunity and auto-immunity in modern humans. One of the most interesting results was a putative introgressed Neanderthal SNP in interleukin 18 (*IL18*), a gene with a central role in the innate immune response and the development of bacterial sepsis. *IL18* expression is induced by products of both gram-positive and gram-negative bacteria. There is evidence for antagonistic pleiotropy in the role of *IL18* in human health and disease: IL18 induces interferon gamma, which can protect against infection; but increased IL18 cytokine signalling is also associated with allergic reaction and development of sepsis^35^. The introgressed *IL18* SNP rs1834481 is associated with decreased serum IL18 levels. If Neanderthals were particularly at risk from bacterial sepsis, this could have created a selection pressure for reduced *IL18* expression^36^.

A Neanderthal allele was also identified in *TNPO3*^7^, a gene associated with increased risk of systemic lupus erythematosus. There is some evidence to suggest that SLE may be triggered by an aberrant response to infection^37^. Further SNPs were identified which play a role in Crohn’s disease, both to increase and decrease susceptibility to this auto-immune disease. A separate study of the same Altai genome by Vernot and Akey^6^ identified a Neanderthal variant of *RNF34* in modern Asian and European genomes, a ring-finger protein with anti-apoptotic functions that interacts with tumour necrosis factor.

There are regions of the genome in which Neanderthal DNA does not persist, seemingly removed by purifying selection for disadvantageous phenotypes; the continued presence of genetic variants associated with immunity in some European and Asian genomes suggests that some Neanderthal haplotypes conferred a selective advantage to *Homo sapiens* during the colonisation of Europe and East Asia. However, individual studies of Neanderthal-human admixture use different methods to identify introgressed DNA, and subsequently identify different regions of the human genome as Neanderthal-derived. It is unclear whether this methodological diversity is a strength or a weakness of the field, as the false-positive rate is unknown.

It is also important to note that our interpretation of the function of these genetic variants, and our identification of immunity-related variants, replies upon our knowledge of the function of genes and polymorphisms within the human genome, which is incomplete – for example, there may be many more polymorphisms affecting susceptibility to viral, bacterial or fungal infection which we have not yet identified in modern humans, and therefore cannot identify in Neanderthal genetic data.

## Ancient pathogen genomics

The work of Johannes Krause^38^ and others^39,40^ raises the tantalising possibility of being able to directly test ancient remains for evidence of infection by amplifying the DNA or RNA of the pathogens which infected them in life. As the horizon for amplifying ancient host DNA moves further back in time (most recently, the 400,000 year old mtDNA sequence from Sima de los Huesos in Spain^41^), it is likely that amplifying ancient pathogen DNA from selected skeletal remains of Neanderthals and Denisovans will become possible. Ancient pathogen sequencing is even reaching into the mouths of Mesolithic and early Neolithic individuals, characterising the oral pathogens preserved in dental calculus^42^. Environmental contamination remains a significant issue in studies of ancient pathogen DNA, with careful use of nucleic extraction methods and well-chosen controls necessary for its prevention and identification^43^.

## Infectious disease in the Pleistocene

The paradigm of the first epidemiologic transmission, the hypothesis that epidemic disease did not occur until the transition to agriculture, with larger, denser and more sedentary populations, has been essentially unchallenged since the 1970s. Our views of the infectious disease environment of the Pleistocene period are heavily influenced by skeletal data and studies of contemporary hunter-gatherers^1^. New genetic data – encompassing both hosts and pathogens – has the power to transform our view of the infectious disease landscape experienced by Neanderthals in Europe, and the AMH with whom they came into contact. The Pleistocene hominin environment cannot be thought of as free from infectious disease. It seems likely that the first epidemiologic transition, envisaged as part of the package of the Holocene farming lifestyle, may be fundamentally different in pace or scope than has previously been suggested.

**Table 1:**
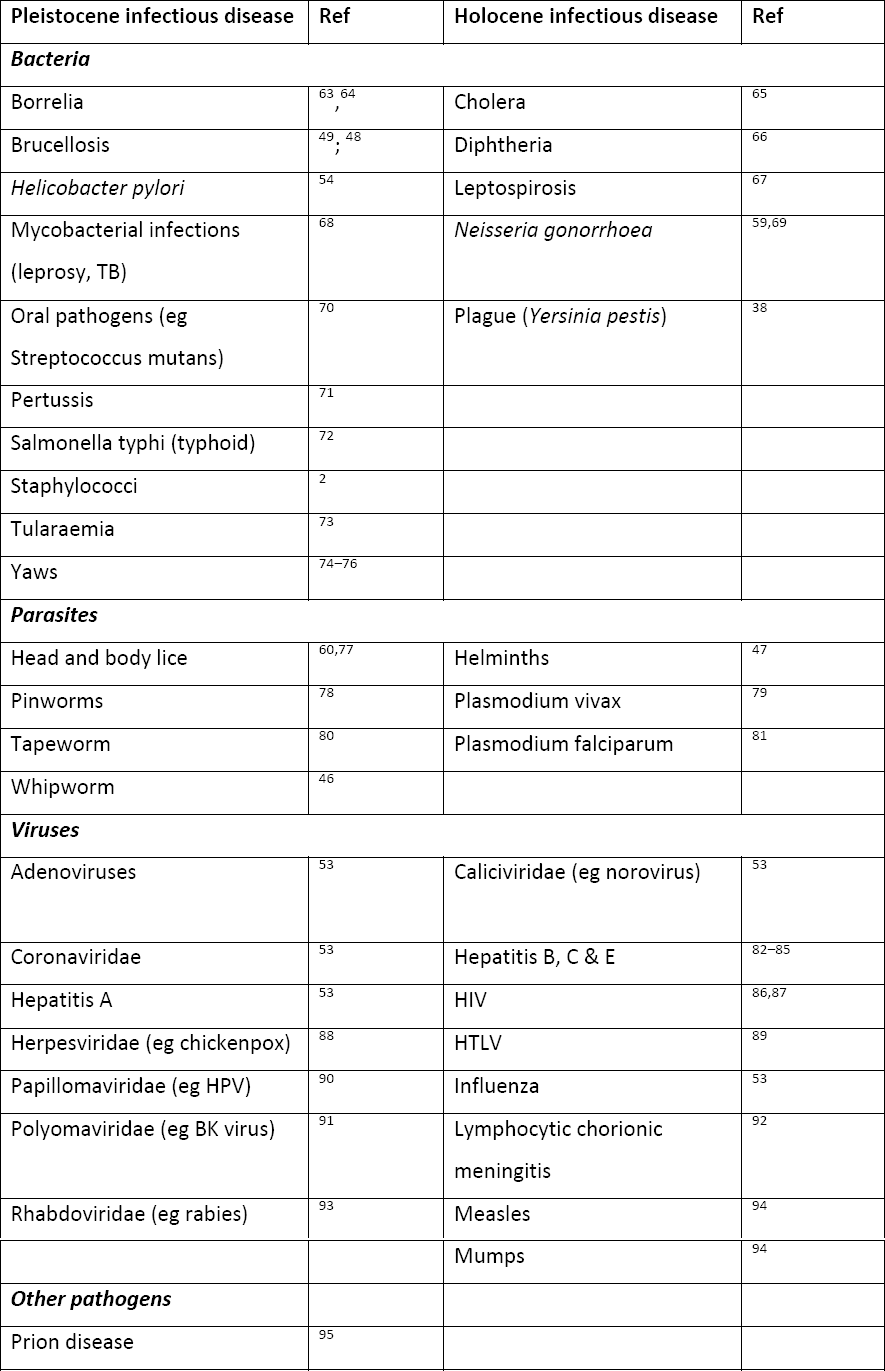
Comparison of European Pleistocene and Early Holocene Disease Packages.

## Challenging the Holocene epidemiologic transition

In the genomes of Neanderthals we can clearly see evidence of the selection pressure exerted by infectious disease. The genome of a 7,000-year-old hunter-gatherer from La Brana in Spain shows similar signals of selection, for example carrying the non-functional form of gene *CASP12* (caspase-12). Functional *CASP12* is associated with an increased risk of bacterial sepsis, and the non-functional form is at or approaching fixation in non-African populations^44^. When considered alongside the reduced expression of Neanderthal *IL18* SNP found in some Europeans and Asians, it is clear that bacterial sepsis was a significant selection pressure on archaic and AMH, long before the assumed arrival of zoonoses with the rise of agriculture in the Holocene.

Sequencing of the Neanderthal and Denisovan genomes has revealed a number of regions of putatively introgressed archaic DNA in modern European and Asian genomes, as discussed above, providing evidence that Neanderthals - and potentially other non-human hominins - experienced significant selection pressure to adapt to infectious disease; these same stretches of DNA may have been advantageous in protecting admixed AMH against the same pathogens.

Paleogenomics provide us with a counterpoint to the AMH skeletal evidence of increasing infectious disease in the Holocene, contributing to a view of Pleistocene Europe as riddled with infectious diseases and parasites. Studying the phylogenetic relationships of extant pathogens has led researchers to conclude that many infectious diseases have been co-evolving with humans and our ancestors for tens of thousands to millions of years. Furthermore, pathogens that were traditionally thought to be zoonoses acquired from herd animals may in fact be anthroponoses, pathogens humans passed to their animals during the rise of agriculture.

It is useful to consider which infectious diseases European Neanderthal populations may have experienced (Table 1). Pleistocene diseases include pathogens which are found in all primates, and are therefore likely to have co-speciated with Neanderthals (also known as heirloom pathogens); and also those pathogens that phylogenetic evidence suggest predate the Holocene, and are therefore potential Neanderthal pathogens. The same infectious diseases would have affected the first AMH in Europe. They are compared to the diseases associated with the transition to agriculture in the Holocene.

Certain pathogens are of particular interest to those studying infectious disease in Neanderthals (see Table 1). Kuhn and colleagues^45^ speculate that a Pleistocene European rock shelter shows evidence of bedding being burned to eliminate parasites and pests. If Pleistocene European AMH were subject to parasites contaminating their bedding, Neanderthals must have been similarly burdened. There are significant tapeworm reservoirs in African primates that have ancient divergence dates from other species^46^ and both Neanderthals and AMH were likely to have carried these parasites. The extent to which they would have caused symptomatic disease is less clear: helminths are often thought to have been a significant source of infectious disease for early foragers, but modern subsistence farmers have higher helminth loads than modern foragers (with the caveat that modern hunter-gatherers/foragers and farmers are not a time capsule of the Pleistocene or Holocene disease landscape^47^).

*Brucella* may be a very ancient human pathogen, despite its modern associations with milk and pastoralism. Phylogenetic analysis of the *Brucella* genus suggests that the different species of *Brucella* diverged tens of thousands of years before the origin of pastoralism and has likely been endemic in wild animal populations for 80,000 – 300,000 years^48^. Brucellosis could therefore have been a disease of Neanderthals and AMH. There are skeletal reports of brucellosis in *Australopithecus africanus,* an order of magnitude earlier than the above estimates^49^. Oral pathogens would also have been a hazard for Neanderthals, not just Holocene farmers. There are reports of dental caries from *Homo heidelbergensis*^50^. Sequencing of Pleistocene dental calculus from AMH and Neanderthals would help researchers to understand the evolution of oral microenvironments during the Pleistocene.

It is therefore likely that Neanderthals were subject to a wide variety of infectious diseases, many of which do not leave skeletal lesions. These pathogens would have had the capacity to cause morbidity and mortality in a variety of settings: infections of dental carries and flesh wounds; childhood diseases (e.g. varicella zoster - chicken pox); gastrointestinal infections; sexually transmitted infections; progressive infections such as leprosy; and many chronic infections which would have been carried for life and only become symptomatic when other infections led to immune suppression, such as tuberculosis and hepatitis.

## Disease exchange

There is as yet no evidence of infectious disease transmission between AMH and Neanderthals, but when considered in the light of the temporal and geographical overlap between the two species^51^ and the evidence of admixture, it must have occurred. There is compelling evidence from Africa of pathogen exchange between humans and other hominins, preserved in the genome of Kaposi’s sarcoma herpes virus (human herpesvirus 8). The K15 gene of KSHV has three highly divergent forms, P, M and N. P is most common, M is found at low frequencies worldwide, and N is rare and found solely in southern Africa^52^. It is thought that the highly divergent M and N forms of K15 introgressed into human KSHV strains through recombination with another herpesvirus that has yet to be detected in modern humans. Based on the divergence dates of the different forms of K15, Hayward and Zong suggest that the M form diverged from the P form 200,000 years ago, and the N form 500,000 years ago. The presence of these other K15 gene forms has arisen through contact with other hominin species who carried their own KSHV-like viruses which speciated with each hominin group. It was originally speculated that the M form of K15 may have originated in a Neanderthal herpesvirus^53^, but the detection of the M form in Africa - where Neanderthal DNA is not detected in living humans - suggests that there would have been one or more unknown hominin species who had contact with AMH in Africa and exchanged pathogen DNA with them. In a sense, the KSHV genome is a fossil record, preserving evidence of past pathogenic interactions between hominins.

*Helicobacter pylori* may be a pathogen which humans transmitted to Neanderthals. *H. pylori* made the out-of-Africa migration with modern humans, estimated to have first infected humans in Africa 88-116kya, and arriving in Europe after 52kya^54^. Chimpanzees do not harbour *H. pylori*, and there is evidence that some African hunter-gatherer groups, such as the Baka, did not acquire *H. pylori* until the last several hundred years, through contact with other groups^55^. The same process of pathogen transmission may have occurred between Neanderthals and AMH.

## Primates, hominins and zoonoses

The close genetic relatedness of AMH and other hominins would only have made it easier for pathogens to jump from one species to another. In the Holocene, wild non-human primates have been the source of acute and chronic infectious diseases which have caused significant mortality: HIV, human T lymphotropic viruses (HTLVs), and vivax and falciparum malaria, for example^56–60^. This demonstrates the ability of infectious diseases to spread between species, through horizontal, vertical or vector-driven disease transmission routes. Humans migrating out of Africa would have been a significant reservoir of tropical diseases, not all of which require vectors for transmission. Likewise, the native Neanderthal populations of Eurasia would have carried hominin-adapted local microbes and parasites.

## Inbreeding depression and Neanderthal immunity

The complete genome sequence of a Neanderthal from the Altai mountains (dated 50,000 B.P.) has also revealed a factor important to our understanding of infectious disease in Neanderthals: inbreeding. The parents of the Altai Neanderthal were as closely related as half-siblings or other similar relationships (e.g. double first cousins). This Neanderthal came from a small effective population, where genetic heterozygosity was low^5^. Infectious diseases would have become an increasingly important factor in Neanderthal mortality, as genetic variants increasing susceptibility to infection became more common in the population and the likelihood of infants being born with primary immune deficiencies increased^61^.

## Conclusion

Analysing the genomes of archaic hominins provides evidence of pathogens acting as a population-level selection pressure, causing changes in genomes that were passed on to descendants and preserved in the genomes of modern Eurasians. Through sequencing ancient pathogen DNA, excavating fossilised parasites, and evidence that Neanderthals had genetic immunity to certain infectious diseases, we will be able to detect pathogens which were previously ‘invisible’ to paleopathology^62^. Skeletal evidence is no longer the sole source of evidence of individual or group-level pathology. Studying genetic data (from host and pathogen) may also point towards new skeletal markers of infection. Comparison of skeletal remains from hominins and hunter-gatherers from the geographical range of the Neanderthals may identify infectious diseases which exerted a significant selection pressure on the Neanderthal genome, and provide evidence of selection on appropriate genetic pathways within the growing collection of ancient human, Neanderthal and Denisovan genomes.

Paleogenomic data must inform our model of the first epidemiologic transition. The view of the Pleistocene infectious disease landscape is being radically altered by analysis of modern and ancient human genomes. Pleistocene hominins were under considerable selection pressure due to infectious disease and the fingerprints of this selection are preserved in ancient and modern genomes. Selection pressure on Neanderthals and on AMH colonising the Neanderthal range must have been maintained over a period of time, or the mutations would have been lost from the modern human gene pool. Clearly, the Pleistocene in temperate regions was not free from acute or chronic infectious diseases and the selection pressures they exert.

Omran^3^ considers parasitic diseases, tuberculosis, pneumonia (respiratory infection) and diarrhoeal diseases to be hallmarks of disease in the early agricultural era of the Holocene, dubbed “the age of pestilence and famine". Anthropological and epidemiological data suggest that many acute infections require large, sedentary populations to be maintained, or an available pool of pastoral animals to act as intermediate hosts^4^, precluding the spread of many infectious diseases in the Pleistocene. In contrast, host and pathogen genetic data suggest a hypothesis of acute respiratory, soft tissue and diarrhoeal diseases having a pre-Holocene association with AMH and Neanderthals. Many of the pathogens thought to have originated in pastoral animals actually originated in humans, including tuberculosis, brucellosis, *Bordetella pertussis*, typhus and typhoid. Subsequently, a number of these infections have become anthroponoses, infections that humans have passed to ruminants and poultry during the transition to agriculture and the intensification of farming. Many of the infectious diseases previously thought to be hallmarks of the first epidemiologic transition (placed in the Holocene) have their roots in the Pleistocene. The rise of agriculture during the Holocene may have intensified their impact on modern human health and changed disease transmission dynamics, but it no longer makes sense to think of the transition as a change from health to pestilence. Infectious disease was entrenched in the Pleistocene landscape, and was an evident selection pressure for hominins in temperate and tropical latitudes. For the Neanderthal population of Eurasia, exposure to new human pathogens carried out of Africa may have been catastrophic.

The model of the first epidemiologic transition must continually develop to include new genetic data. We must also consider whether the first epidemiologic transition was a much longer process than previously envisaged, spanning both the Pleistocene and early Holocene epochs and several hominin species, not just *Homo sapiens.* The analysis of ancient genomes demonstrates that human behavioural patterns (in this case a shift to agricultural subsistence) should not be used as an ecological proxy to explain shifting trends in the co-evolutionary relationship between pathogens and human populations.

## Acknowledgements

The authors would like to thank Professor Robert Foley for reading a draft of this paper.

